# Disruption of the gut microbiota attenuates epithelial ovarian cancer sensitivity to cisplatin therapy

**DOI:** 10.1101/2020.06.16.155226

**Authors:** Laura M. Chambers, Emily L. Esakov, Chad Braley, Lexie Trestan, Zahraa Alali, Defne Bayik, Justin D. Lathia, Naseer Sangwan, Peter Bazeley, Amy S. Joehlin-Price, Mohammed Dwidar, Adeline Hajjar, Philip P. Ahern, Jan Claesen, Peter Rose, Roberto Vargas, J. Mark Brown, Chad Michener, Ofer Reizes

## Abstract

Epithelial Ovarian Cancer (EOC) is the leading cause of gynecologic cancer death. Despite many patients achieving remission with first-line therapy, up to 80% of patients will recur and require additional treatment. Retrospective clinical analysis of OC patients indicates antibiotic use during chemotherapy treatment is associated with poor overall survival. We assessed whether antibiotic (ABX) therapy would impact growth of EOC and sensitivity to cisplatin in murine models. Immune competent or compromised mice were given control or ABX containing water (metronidazole, ampicillin, vancomycin, and neomycin) before being intraperitoneally injected with murine EOC cells. Stool was collected to confirm microbiome disruption and tumors were monitored, and cisplatin therapy was administered weekly until endpoint. EOC tumor-bearing mice demonstrate accelerated tumor growth and resistance to cisplatin therapy in ABX treated compared with nonABX treatment. Stool analysis indicated most gut microbial species were disrupted by ABX treatment except for ABX resistant bacteria. To test for role of the gut microbiome, cecal microbiome transplants (CMTs) of microbiota derived from ABX or nonABX treated mice were used to **r**ecolonize the microbiome of ABX treated mice. nonABX cecal microbiome was sufficient to ameliorate the chemoresistance and survival of ABX treated mice indicative of a gut derived tumor suppressor. Mechanistically, tumors from ABX treated compared to nonABX treated mice contained a high frequency of cancer stem cells that were augmented by cisplatin. These studies indicate an intact microbiome provides a gut derived tumor suppressor and maintains chemosensitivity that is disrupted by ABX treatment.

**Significance:** Platinum resistance is associated with poor prognosis and reduced therapeutic options for ovarian cancer patients. We identifed a tumor suppressive role of the gut microbiome that is disrupted upon antibiotic therapy.

## Background

Epithelial ovarian, peritoneal and fallopian tube carcinomas (EOC) are a leading cause of gynecologic cancer related death (1). Following EOC diagnosis, patients are treated with a combination of cytoreductive surgery and platinum-taxane based chemotherapy (2–8). Unfortunately, despite many women achieving remission with first-line therapy, up to 80% of patients will recur and require additional treatment (2–8). After disease recurrence, the interval from last treatment with platinum chemotherapy has important therapeutic and prognostic implications (9). Patients with platinum resistant EOC have fewer treatment options, reduced response rates to chemotherapy and poor prognosis compared to those with platinum sensitive disease (9). There is a significant unmet need to further understand the mechanisms of platinum resistance in EOC and to advance current therapeutic options for these patients.

The gut microbiome has many roles in maintenance of human health, and has been increasingly linked with many disease states, including cancer (10–18). Recent evidence suggests that the gut microbiome may modulate responses to cancer treatment, including traditional chemotherapy and immunotherapy (10–19). In a study by Routy et al, resistance to immune checkpoint inhibitors was linked to abnormal gut microbiome composition following treatment with antibiotics (18). Similarly, Iida et al. evaluated the role of the gut microbiome upon platinum chemotherapy response in mice with lymphoma, colon cancer and melanoma tumors (14). The impact of oxaliplatin treatment was attenuated in mice treated with antibiotics showing decreased tumor regression and worse survival, compared to non-antibiotic control animals.

Antibiotic therapy is frequently used during cancer treatments for both prophylaxis and treatment of infections. Studies have demonstrated that receipt of antibiotics during both systemic chemotherapy and immunotherapy negatively impacts oncologic outcomes. In a study of patients with relapsed lymphoma and leukemia receiving either cisplatin or cyclophosphamide on two clinical trials, those that received antibiotics against gram positive bacteria during platinum chemotherapy had a significant reduction in overall treatment response, time to recurrence and overall survival (17). Among patients with epithelial ovarian cancer, studies have demonstrated that treatment for surgical site infection following primary cytoreductive surgery is associated with worse survival (20). In a recent retrospective analysis, EOC patients who received antibiotics, primarily against gram positive bacteria, during primary platinum chemotherapy had reduced overall (>17 months) and progression free survival (>24 months) compared to patients who received no antibiotic intervention (21). These studies suggest that an intact gut microbiome provides a protective microenvironment, and disruption leads to accelerated tumor growth and chemotherapy resistance, including platinum agents. Here, we investigated the impact of antibiotic mediated disruption of the gut microbiome on EOC tumor growth and platinum chemotherapy response in pre-clinical models of EOC.

## Methods

### Cell Lines

ID8, ID8-VEGF (syngeneic) and OV81 (human) EOC cell lines were cultured in Dulbecco Modified Eagle Medium (DMEM) media containing heat inactivated 5% FBS (Atlas Biologicals Cat # F-0500-D, Lot F31E18D1) and grown under standard conditions. HEK 293T/17 (ATCC CRL-11268) cells were plated and co-transfected with Lipofectamine 3000 (L3000015 Invitrogen), 3rd generation packaging vectors pRSV-REV #12253, pMDG.2 #12259, and pMDLg/pRRE #12251 (Addgene) and lentiviral vector directing expression of luciferase reporter pHIV-Luciferase #21375 4.5 μg (Addgene). Viral particles were harvested, filtered through a 0.45 μm Durapore PVDF Membrane (Millipore SE1M003M00) and added to cell line culture media. Viral infections were carried out over 72 hours and transduced cells were selected by their resistance to 2 μg/mL puromycin (MP Biomedicals 0219453910).

### Animal Studies

Female C57BL/6J (BL/6) mice were purchased from Jackson Laboratories (Bar Harbor, ME) at 6 weeks of age. Female NOD.Cg-Prkdc<scid>Il2rg<tm1Wjl>SzJ (NSG) mice were purchased from the Cleveland Clinic BRU at 6 weeks of age. Experimental animals were housed and handled in accordance with Cleveland Clinic Lerner Research Institute IACUC guidelines. Mice (BL/6 or NSG) were given either control or antibiotic (0.5 g/L vancomycin, 1 g/L neomycin sulfate, 1 g/L metronidazole, 1 g/L ampicillin) (ABX) (Fisher Scientific) containing water for 2 weeks prior to IP injection of ID8-LUC (5×10^6^) ID8-VEGF (5×10^6^) or OV81 (5×10^6^) cells. The ABX containing water has been previously shown to be sufficient to deplete all detectable commensal bacteria (22, 23). Mice remained on either control or ABX containing water for the duration of the study. At 10 weeks of age, mice were treated IP with either cisplatin (0.5mg/kg, Spectrum Chemical or vehicle (PBS) weekly until human endpoint of total tumor burden exceeding 150mm^3^ or debilitating ascites development.

### Murine CMT Studies

Following two weeks of ABX or normal water treatment of mice, necropsies were performed, and ceca harvested and transported to an anaerobic coy chamber for further processing. Ceca were dispersed with anaerobic PBS by vortexing for 10 minutes and set for 10 minutes to allow debris to settle. The top portion was aliquoted into 1mL crimp-top tubes and stored at −80C until used. ABX CMT mice received 1 week of ABX treatment as described above followed by cesation of ABX and 2 doses of CMT material from ABX treated mice through oral gavage and put back on ABX water throughout. H_2_O CMT mice were also treated for 1 week with ABX and received 2 doses of CMT material from H_2_O treated mice through oral gavage. All mice received an IP injection of 5×10^6^ ID8-LUC cells on day 27 of the study and monitored for tumor progression via IVIS imaging as described above as well as pre-determined endpoint criteria of debilitating ascites development or a body composition score (BCS) < 2.

### Murine Body Composition Scoring

While gently restraining a mouse by holding the base of its tail, the observer (blinded to treatment group) used the thumb and index finger of the other hand to palpate the degree of muscle and fat over the sacroiliac region. A score from 1-5 was given to each mouse weekly following IP tumor cell injection based on previous literature (24, 25). Based on established IACUC protocols #2018-2003, a BCS score of 2 or lower was defined as meeting endpoint criteria.

### Tumor Monitoring by Trans abdominal Ultrasound (TAUS)

For the ID8-LUC and ID8-VEGF cohorts, TAUS surveillance was initiated seven days following IP tumor injection as previously described (26). TAUS was performed every 10 days until study endpoint. Mice were anesthetized using isoflurane (DRE Veterinary) and placed in the supine position. Following the removal of abdominal hair using Nair (Church & Dwight Co. Inc.), sterile ultrasound gel was applied to the abdomen. TAUS was performed using Vevo2100 (VisualSonics) using the abdominal imaging package and MS550D probe (40Hz). For each mouse, the abdomen was assessed for tumor in four quadrants. Tumors were noted to be absent or present at each assessment. Tumor length and width were recorded and tumor volume was calculated using the formula: (Length*(Width^2^))/2.

### Tumor Monitoring by 2D IVIS Imaging

For the OV81-LUC and CMT study ID8-LUC cohort of mice, bioluminescence images were taken with IVIS Lumina (PerkinElmer) using D-luciferin as previously described (27). Mice received an IP injection of D-luciferin (Goldbio LUCK-1G, 150mg/kg in 150μL) under inhaled isoflurane anesthesia. Images were analyzed (Living Image Software) and total flux reported in photons/second/cm^2^/steradian for each mouse abdomen. All images were obtained using the automatic exposure feature and mice were imaged individually.

### 16S rDNA Sequencing

At 5 pre-determined time points (baseline, 2 weeks, tumor engraftment, 5 weeks, and endpoint, see **Figure 1A** schematic), stool was collected from each mouse and frozen in micro centrifuge tubes at −80C. Upon completion of the study, 16S rDNA was isolated following the standard protocol and (QIAamp PowerFecalPro Kit) (Qiagen). Isolated samples were sent to Miami University of Ohio on dry ice for 16S V4 rDNA processing. Data analysis was completed by the Microbiome and Composition Analytics Core facility as follows: Individual fastq files without non-biological nucleotides were processed using Divisive Amplicon Denoising Algorithm (DADA) pipeline (28). The output of the dada2 pipeline (feature table of amplicon sequence variants (an ASV table)) was processed for alpha and beta diversity analysis using *phyloseq (29)*, and microbiomeSeq (http://www.github.com/umerijaz/microbiomeSeq) packages in R. Alpha diversity estimates were measured within group categories using estimate_richness function of the *phyloseq* package *(29)*. Multidimensional scaling (MDS, also known as principal coordinate analysis; PCoA) was performed using Bray-Curtis dissimilarity matrix (30) between groups and visualized by using *ggplot2* package (31). We assessed the statistical significance (*P* < 0.05) throughout and whenever necessary, we adjusted *P*-values for multiple comparisons according to the Benjamini and Hochberg method to control False Discovery Rate (32) while performing multiple testing on taxa abundance according to sample categories. We performed an analysis of variance (ANOVA) among sample categories while measuring the of α-diversity measures using plot_anova_diversity function in *microbiomeSeq* package (http://www.github.com/umerijaz/microbiomeSeq). Permutational multivariate analysis of variance (PERMANOVA) with 999 permutations was performed on all principal coordinates obtained during PCoA with the *ordination* function of the *microbiomeSeq* package. Linear regression (parametric test), and Wilcoxon (Non-parametric) test were performed on ASVs abundances against coprostanol levels using their base functions in R (33).

**Figure 1.**
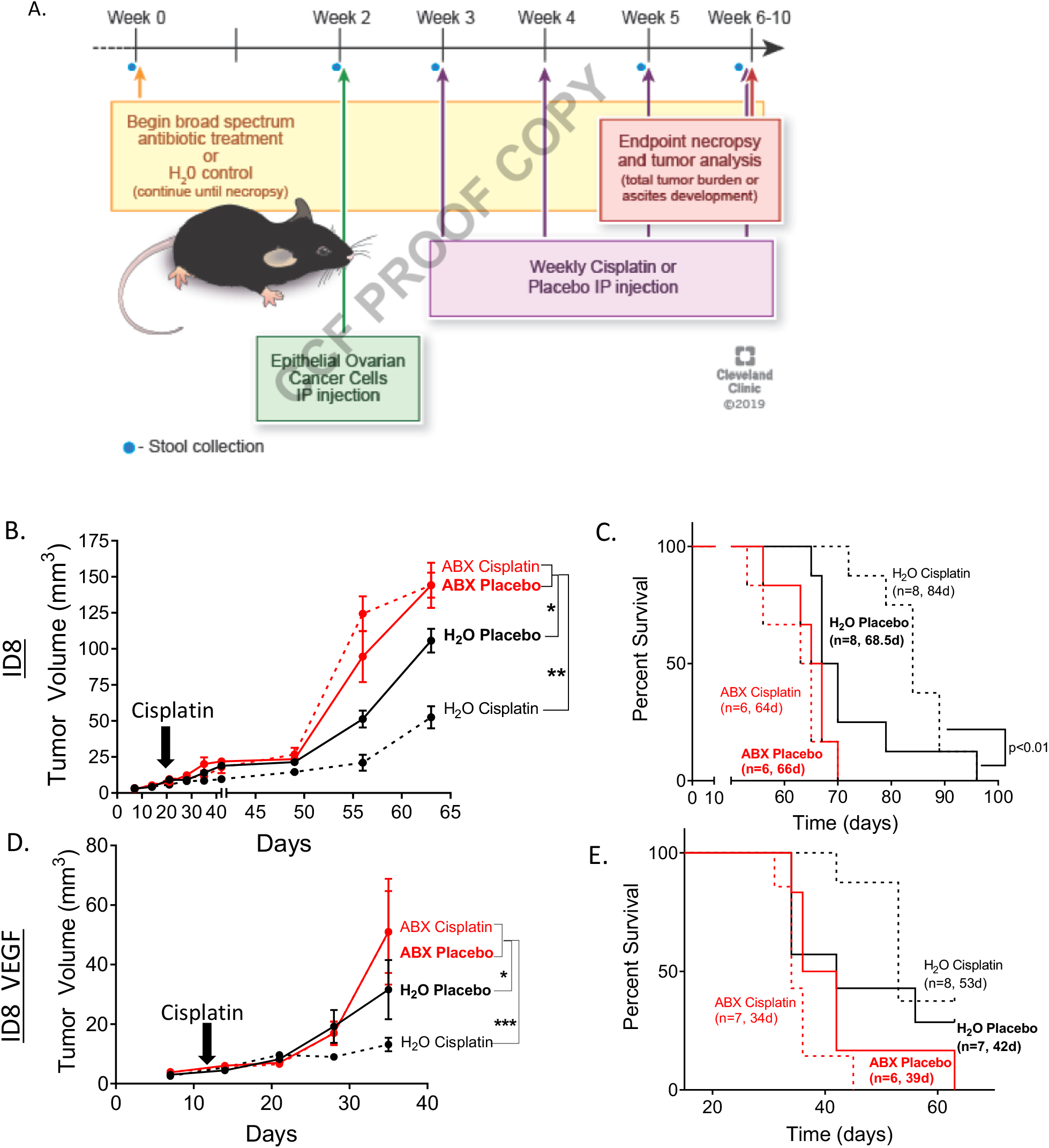
Tumor growth is increased and median survival is decreased in ABX treated Bl/6 mice, regardless of cisplatin therapy. Mice treated with antibiotics following the schematic study design (A) exhibited increased tumor growth regardless of cisplatin therapy following injection of both ID8 (B) and ID8 VEGF (D) cells in C57 BL/6 mice. Antibiotic treated ID8 (C) and ID8 VEGF (E) injected C57 Bl/6 mice (red) had significantly decreased median survival compared to H_2_O treated mice (black). Further, cisplatin therapy (dashed lines) increased H_2_O treated murine survival (black, ID8 p<0.01), but did not alter ABX treated murine survival (red), in either cohort. (n=6 or otherwise denoted in graphs, 2-way ANOVA *p<0.05, **p<0.01, ***p<0.005 survival curve analysis: Gehan-Breslow-Wilcoxon test placebo vs cisplatin).

### Tumor RNA Sequencing

Tumors were collected at endpoint from NSG ID8 mice and snap frozen in liquid nitrogen. Prior to RNA extraction (Takara Nucleospin), tumors were crushed using ice-cold mortar and pestle under sterile conditions. Following RNA isolation, cDNA libraries were prepared by the LRI Genomics Core facility and sent to Macrogen Inc. for RNA sequencing. The RNA sequencing results were analyzed by the Cleveland Clinic Quantitative Health Sciences’ Bioinformatics Consulting Service as follows:

Raw sequence FASTQ files underwent read quality assessment with FASTQC (version 0.11.8, https://www.bioinformatics.babraham.ac.uk/projects/fastqc/) (34). Ribosomal RNA content was evaluated with SortMeRNA (version 2.1) (35). Adapter removal and quality trimming was performed with Trimgalore (version 0.5.0, https://github.com/FelixKrueger/TrimGalore). Adapter removed, trimmed reads were aligned to the Mus musculus genome assembly (GRCm38 version 99, ftp://ftp.ensembl.org/pub/release99/fasta/mus_musculus/dna/Mus_musculus.GRCm38.dna.primary_assembly.fa.gz) using STAR (version 2.6.1d)(36), and duplicates were marked with picard (version 2.18.27, http://broadinstitute.github.io/picard/). Alignments were assessed for quality with qualimap (version 2.2.2)(37) and rseqc (version 3.0.0)(38), and for library complexity saturation with preseq (version 2.0.3)(39). Aligned reads were counted at the exon level and summarized at the gene level with the featureCounts tool from the Subread package (version 1.6.4)(40), using annotations for build GRCm38 (version99, ftp://ftp.ensembl.org/pub/release99/gtf/mus_musculus/Mus_musculus.GRCm38.99.gtf.gz). Normalization and differential expression analysis was performed with the R (version 3.6.3)(41) package DESeq2 (version 1.26.0)(42). Size factor and dispersion estimation were performed with default settings. Comparison estimate p-values for H_2_O-Cisplatin vs. H_2_O-Placebo, Antibiotic-Cisplatin vs. Antibiotic-Placebo, Antibiotic-Cisplatin vs. H_2_O-Cisplatin, and Antibiotic-Placebo vs. H_2_O-Placebo were extracted using multiple testing adjustment with the method of Benjamini and Hochberg (43), and an independent filtering significance cut-off of 0.05 (44). Log2-fold change estimates for each comparison were shrunken with the function lfcShrink from the DESeq2 package, using default settings. The web tool DAVID bioinformatics database was utilized to assess the top 100 significantly expressed genes from each group comparison for enrichment of GO terms (45).

### Flow Cytometry

Upon murine necropsy, ascites, splenocytes, and bone marrow was collected, processed for single cell suspension through filtration and stored in freezing media (10% DMSO in FBS, Atlas Biologicals) at – 80C. All single cells suspensions were thawed on ice, washed once with 2%BSA in PBS and stained with live/dead UV stain (Invitrogen) and then blocked in FACS buffer (PBS, 2% BSA) containing FcR blocking reagent at 1:50 (Miltenyi Biotec) for 15 minutes. After live/dead staining and blocking, antibody cocktails for myeloid or lymphoid panels (**Supplemental Table 1**), were incubated with cells at a 1:50 dilution for 20 minutes on ice before being washed and suspended in FACS buffer for analysis. Cell populations were analyzed using an LSRFortessa (BD Biosciences), and populations were separated and quantified using FlowJo software (Tree Star Inc.). Gating methods for myeloid and lymphoid populations were performed following standardized gating strategies previously described (46, 47). For a complete gating strategies please see **Supplemental Figure 1**.

### Tumorsphere Formation Studies

Upon murine necropsy, Total tumor collected per mouse was dissociated using standard methods with a Papain dissociation kit (Worthington Biologicals). Following filtration through a 40 micron filter, single cells were counted cultured in serial dilutions in a non-adherent 96-well plate (Sarstedt) with 200 μl serum-free DMEM/F12 medium supplemented with 20 ng/mL basic fibroblast growth factor (Invitrogen), 10 ng/mL epidermal growth factor (BioSource), and 2% B27 (vol/vol) (Invitrogen). Tumorsphere-formation was scored following 2 weeks incubation using a phase contrast microscope. The sphere initiation frequency was calculated using an extreme limiting dilution algorithm (ELDA) (http://bioinf.wehi.edu.au/software/elda/).

### In vitro ABX cisplatin sensitivity studies

ID8-LUC and ID8-VEGF cells were plated at 1,000 cells per well and treated with ampicillin, vancomycin, metrinodazole, and neomycin in combinations of the same ratios used in the murine studies for 72 hours, wherein proliferation was measured using Cell Titre Glo (Promega) and compared to vehicle treated control. Additionally following incubation for 7 days with ampicillin, vancomycin, metrinodazole, and neomycin the IC50 of cisplatin (Spectrum Chemical) was assessed compared to vehicle pretreated controls also utilizing Cell Titre Glo (Promega).

### Statistics

All data are presented as mean +/− standard error of the mean. All statistical analysis was performed in GraphPad Prism v8 unless otherwise noted. Replicate numbers and p-values are presented in figure legends.

### Study Approvals

All murine studies were completed in accordance with the Institutional Animal Care and Use Committee guidelines, approval # 2018-2003. All studies utilizing lenti-viral particle generation were completed in accordance with the Institutional Biosafety Committee guidelines, approval #IBC0920.

## Results

### EOC tumors exhibit accelerated growth and attenuated sensitivity to cisplatin in antibiotic treated mice

We tested the hypothesis that an antibiotic driven disruption of the microbiome impacts tumor development and chemotherapy sensitivity in EOC. To this end, the following study paradigm (**Figure 1A**) was implemented wherein C57 BL/6J (BL/6) female mice at 6 weeks of age were provided water or antibiotic containing water (ABX; ampicillin, neomycin, metronidazole, and vancomycin) (23, 48), which was sustained for the duration of the study. After 2 weeks, mice were injected intraperitoneally (IP) with murine EOC lines ID8 or ID8-VEGF, syngeneic with BL/6 mice, and tumor growth was monitored by transabdominal ultrasound (TAUS) weekly for the course of the study (26). ID8 and ID8-VEGF cell lines were utilized as they are highly characterized and closely recapitulate human ovarian cancer progression. The ID8-VEGF cell line is slightly more severe causing an increase in angiogenesis and ascites development (49, 50). At 2 weeks post tumor cell injection, mice were injected IP with cisplatin or vehicle weekly for the remainder of the study. Mice met study endpoint at tumor burden of >150mm^3^ or humane endpoint, including debilitating ascites development. Upon necropsy, EOC tumor phenotype was confirmed through histological assessment of H&E stained tissue sections as well as benign adjacent omentum (data not shown). Our findings indicate that EOC tumor growth is significantly increased by ABX treated mice compared to controls. In the presence of ABX, cisplatin therapy had attenuated efficacy compared to in the control treated mice indicating development of cisplatin resistance (**Figure 1 B, D**). As expected from the tumor burden, median survival was decreased in the ABX treatment groups compared to controls. The ID8 cohort exhibited a median survival of 66 and 64 days in the ABX placebo and cisplatin groups compared to 68.5 and 84 days in the H_2_O placebo and cisplatin groups, respectively **(Figure 1 C).** This result was paralleled in the ID8-VEGF cohort of mice with a median survival of 39 and 34 days in the ABX placebo and cisplatin groups compared to 42 and 53 days in the H_2_O placebo and cisplatin groups respectively **(Figure 1 E)**. The ID8-VEGF mice reached the endpoint earlier than the ID8 cohort secondary to large volume ascites development.

### Limited impact on immune populations in ascites of EOC by broad spectrum antibiotics

The majority of EOC patients present with advanced disease (stage III or stage IV) that may include ascites (51, 52). Although the underlying mechanisms of ascites development are largely unknown, previous studies have reported that immune cells such as NK cells and macrophages found in peritoneal ascites may play a key role in tumor cell invasiveness and growth (53–56). To this end, we analyzed the immune cell populations within the peritoneal ascites fluid through flow cytometry. Our flow cytometry evaluation of immune cell populations within the peritoneal ascites was designed to focus on both myeloid and lymphoid populations (see **Supplemental Table 1 and Figure 1** for antibody panels and gating strategies). Our findings indicated no significant difference in myeloid (**Supplemental Figure 2A**) or lymphoid (**Supplemental Figure 2B**) cell populations in the ascites of mice following ABX therapy in the presence or absence of cisplatin therapy in the BL/6 ID8-VEGF cohort. Additionally, no significant differences were observed between myeloid or lymphoid cell populations within collected splenocytes **(Supplemental Figure 3A, B)** or bone marrow **(Supplemental Figure 3C, D)** at endpoint.

**Figure 2.**
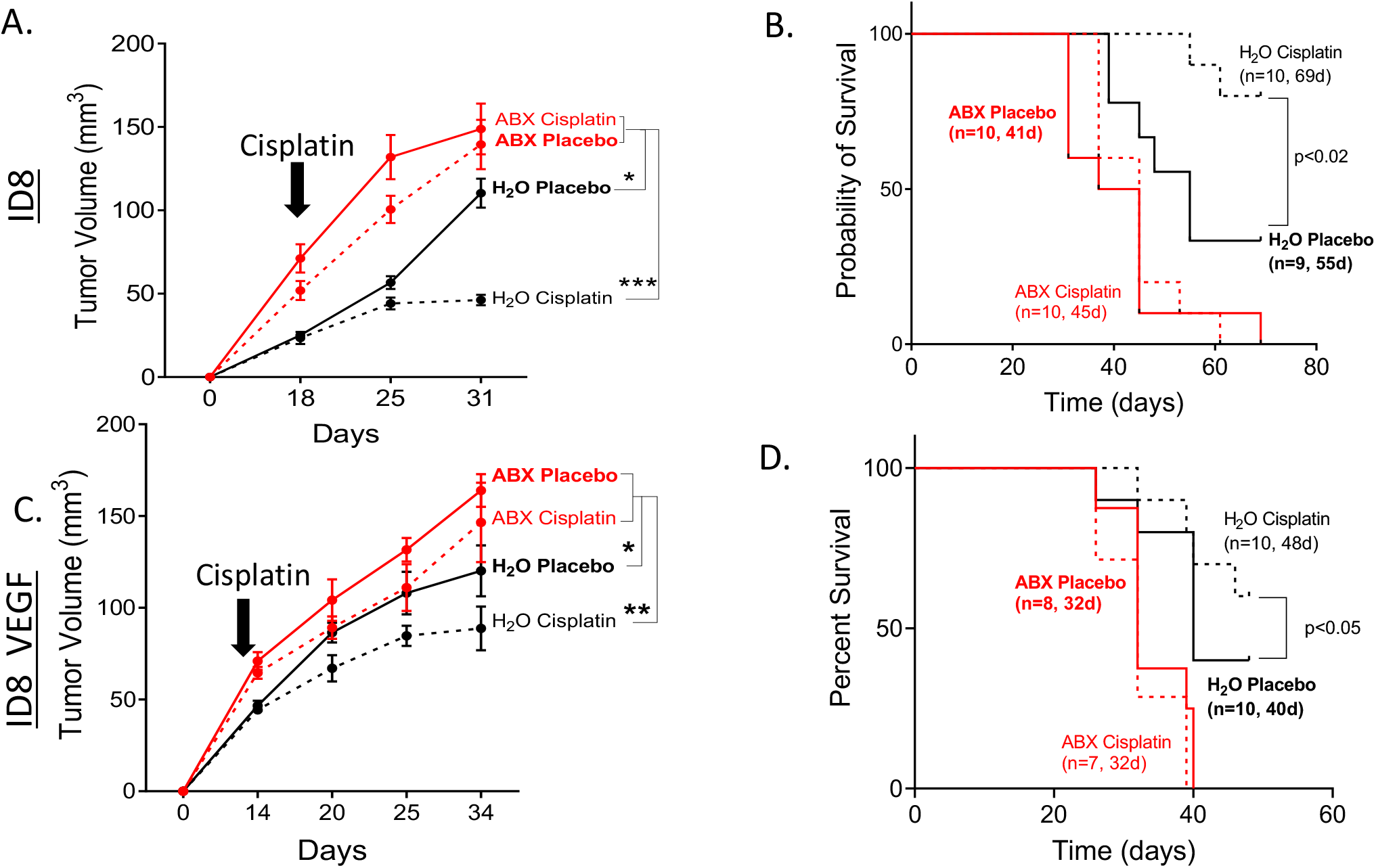
Tumor growth is increased and median survival is decreased in ABX treated NSG mice, regardless of cisplatin therapy. Mice treated with antibiotics exhibited increased tumor growth regardless of cisplatin therapy following injection of both ID8 (A) and ID8 VEGF (C) cells in C57 BL/6 mice. Antibiotic treated ID8 (B) and ID8 VEGF (D) injected NSG mice (red) had significantly decreased median survival compared to H_2_O treated mice (black). Further, cisplatin therapy (dashed lines) increased H_2_O treated murine survival (black, ID8 p<0.01), but did not alter ABX treated murine survival (red), in either cohort. (n=6 or otherwise denoted in graphs, 2-way ANOVA, *p<0.05, **p<0.01, p<0.005 survival curve analysis: Gehan-Breslow-Wilcoxon test placebo vs cisplatin).

**Figure 3.**
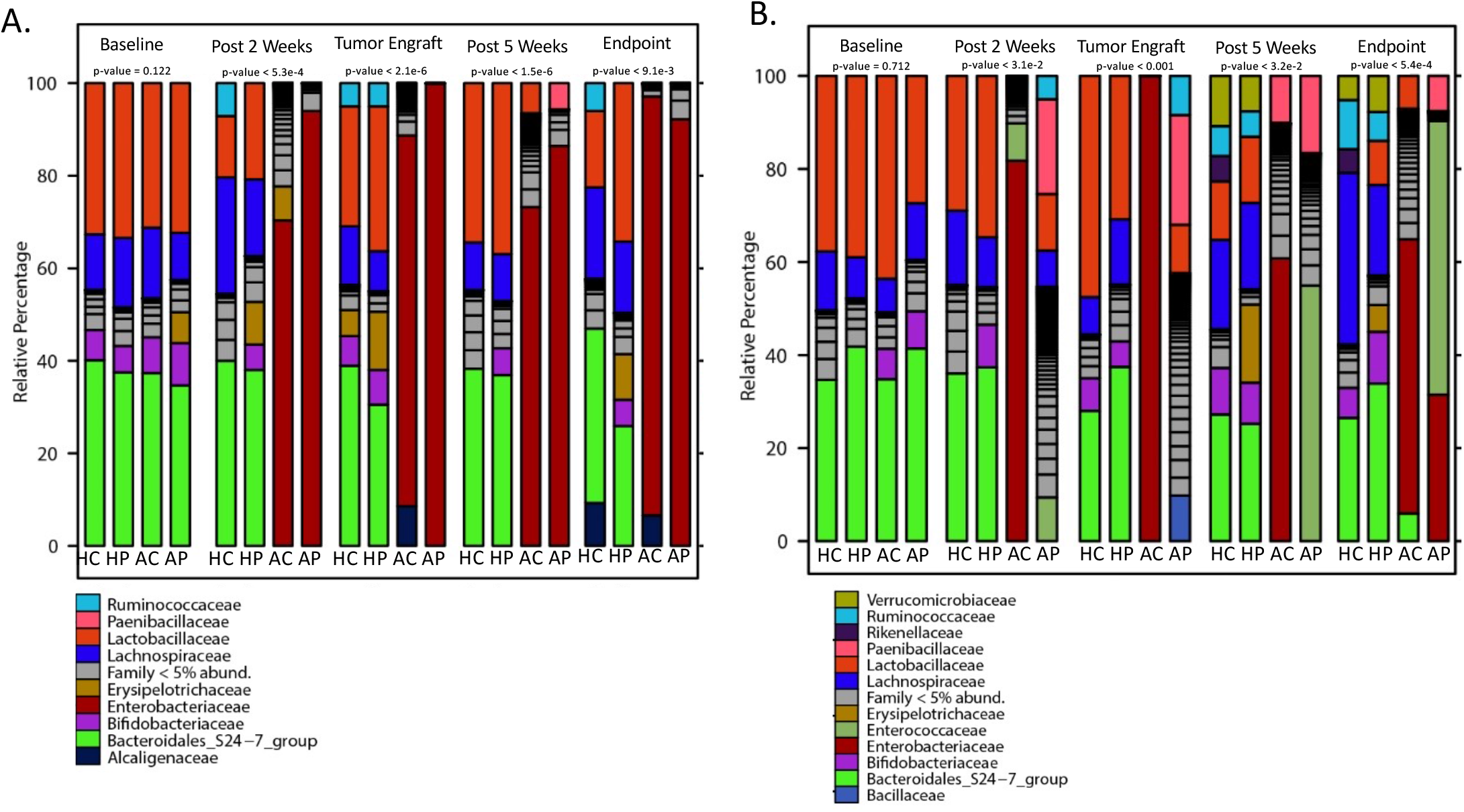
ABX therapy results in microbial alterations over time in the gut microbiome of C57Bl/6 mice. Specific strains of bacteria are altered in the ID8 (A) and ID8 VEGF (B) cohorts of C57 Bl/6 mice following antibiotic treatment over time. n= 8 mice per group (HC: H_2_O Cisplatin, HP: H_2_O Placebo, AC: ABX Cisplatin, AP: ABX Placebo) n=8 mice per group per time point.

### Disruption of the immune system does not significantly impact the accelerated EOC tumor growth or reduced cisplatin sensitivity in antibiotic treated mice

To assess whether the observed augmentation of tumor phenotype and cisplatin resistance following ABX treatment is dependent on an intact immune system, the same study paradigm was utilized in NOD.Cg-Prkdc<scid>Il2rg<tm1Wjl>SzJ (NSG) immuno-deficient mice. The NSG cohort treated with ABX displayed accelerated tumor growth with more rapid onset compared to the BL/6 cohort, with no benefit of cisplatin (**Figure 2 A, C**). The time to tumor progression was accelerated and the effect on median survival was even more significant with the ID8 cohort exhibiting a median survival of 41 and 45 days in the ABX placebo and cisplatin groups compared to 55 and 69 days in the H_2_O placebo and cisplatin groups respectively (**Figure 2 B**). A similar phenotype was observed in the ID8-VEGF cohort of NSG mice (**Figure 2 D**). Additionally, the peritoneal ascites fluid at endpoint was analyzed by flow cytometry. NSG mice have immature T cells, DCs and macrophages, but functional neutrophils (57). NSG ID8-VEGF cohort ascites exhibited no significant alterations in myeloid (**Supplemental Figure 2 C**) populations in ABX treated mice, regardless of cisplatin therapy compared to H_2_O controls.

### ABX treatment results in comparable disruptions of the gut microbiome in BL/6 and NSG EOC tumor bearing mice

The ABX treatment regimen we applied was previously shown to be sufficient to deplete detectable commensal bacteria (22, 23). However, to define the response of the gut microbiome to broad spectrum antibiotic therapy, stool was collected at baseline, at 2 weeks, at tumor engraftment, at 5 weeks and at endpoint necropsy and then processed for 16S rRNA gene sequencing. The microbiome was analyzed from the following cohorts: BL/6 ID8, BL/6 ID8-VEGF, NSG ID8, and NSG ID8-VEGF.

#### C57 BL/6 cohort

In total, 2,801,471 and 2,411,554 high-quality and usable reads were obtained from fecal samples of 5 mice per treatment group in the ID8 and ID8-VEGF BL/6 cohorts from sequencing the 16S rRNA gene, with an average length of 210 base pairs (bps). There were 3,184 amplified sequence variants (ASVs) in all ID8 and ID8-VEGF BL/6 samples. Pairwise comparisons within both the ID8 and ID8-VEGF cohorts revealed statistically significant differences in alpha diversity between temporal collections, as analyzed by the Shannon diversity index (ANOVA) **(Supplemental Figure 4 A, B)**. Additionally, the Bray-Curtis dissimilarity based beta diversity comparisons between collection time points in the ID8 and ID8-VEGF cohorts were significant at p= 0.003 and p=0.001 respectively **(Supplemental Figure 4 C, D).**

**Figure 4.**
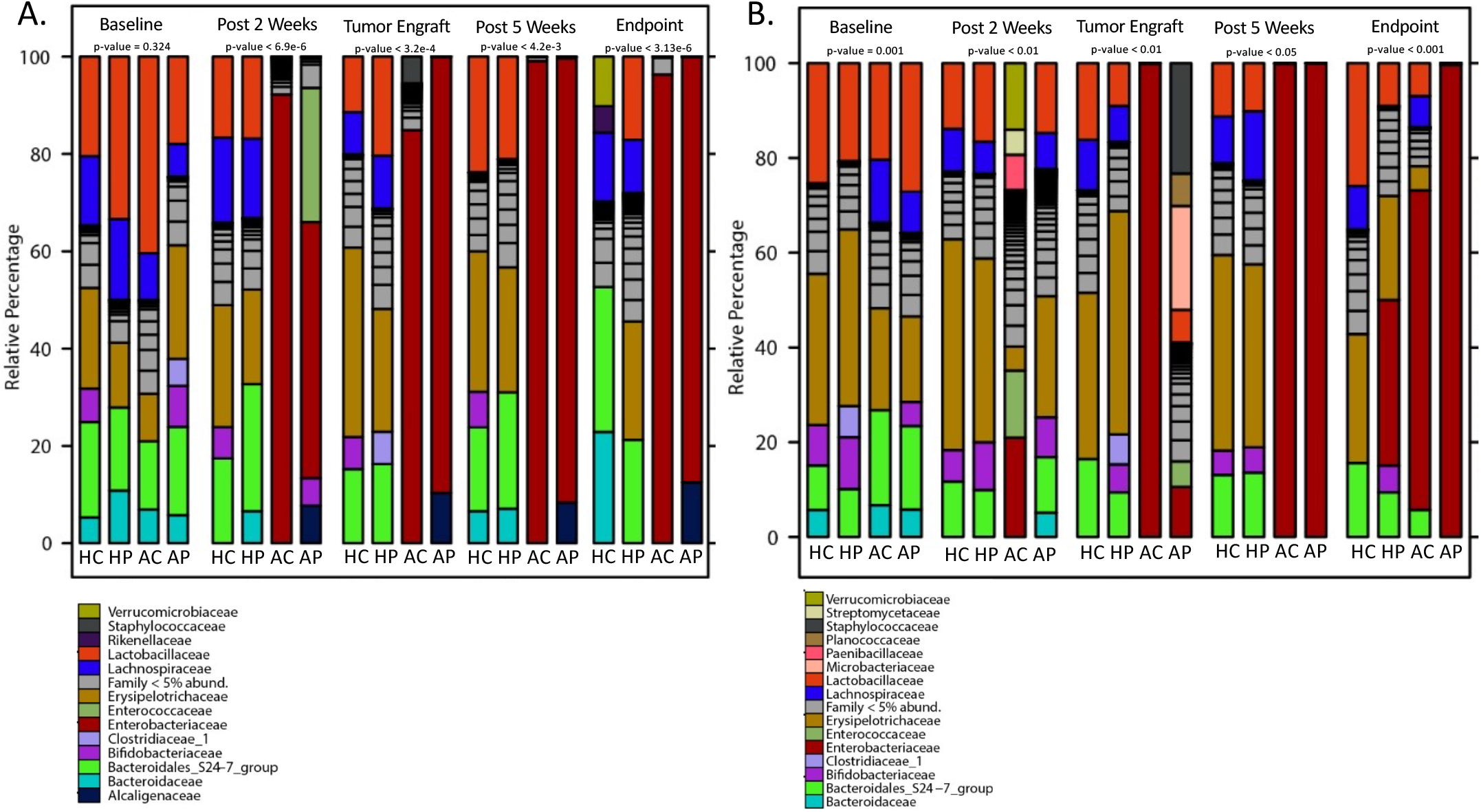
ABX therapy results in microbial alterations over time in the gut microbiome of NSG mice. Specific strains of bacteria are altered in the ID8 (A) and ID8 VEGF (B) cohorts of NSG mice following antibiotic treatment over time. n= 8 mice per group (HC: H_2_O Cisplatin, HP: H_2_O Placebo, AC: ABX Cisplatin, AP: ABX Placebo) n= 8 mice per group per time point.

Antibiotic treated groups displayed significantly less diverse fecal microbial communities compared to control water treated groups regardless of cisplatin therapy. Specifically, the relative abundance of some *Proteobacteria* including *Enterobacteriaceae* and *Parasutterella* was increased in the antibiotic treated ID8 groups while the abundance of *Enterobacteriaceae* was increased in the antibiotic treated ID8-VEGF groups (**Figure 3 A, B**). These were significantly increased by 2 weeks post antibiotic therapy in the mice being co-treated with cisplatin but took up to 5 weeks for the mice treated with placebo to obtain the same increase. Being Gram-negative, and mostly facultative anaerobes, the *Proteobacteria* are generally resistant to vancomycin which affects mainly Gram-positive bacteria and are also less susceptible to metronidazole which affects more the anaerobes (58). Furthermore, *Enterobacteriaceae* have multiple antibiotic resistance genes allowing for their survival post ABX therapy (59, 60). Increased abundance of *Parasutterella* has been associated with dysbiosis of the gut microbiome, but the mechanism has not been fully interrogated (61, 62).

#### NSG cohort

In total, 4,784,498 and 2,822,423 high-quality and usable reads were obtained from fecal samples of 5 mice per treatment group in the ID8 and ID8-VEGF NSG cohorts from sequencing the 16S rRNA gene, with an average length 210 bps. There were 6,484 ASVs in all ID8 and ID8-VEGF NSG samples. As with the BL/6 cohort, the pairwise comparisons revealed statistically significant (ANOVA) patterns for alpha diversity between the collection time points in both the ID8 and ID8-VEGF cohorts **(Supplemental Figure 5 A, B)**. Additionally, the Bray-Curtis dissimilarity based beta diversity between collection time points in the ID8 and ID8-VEGF cohorts were both significant at p=0.001 **(Supplemental Figure 5 C, D).**

**Figure 5.**
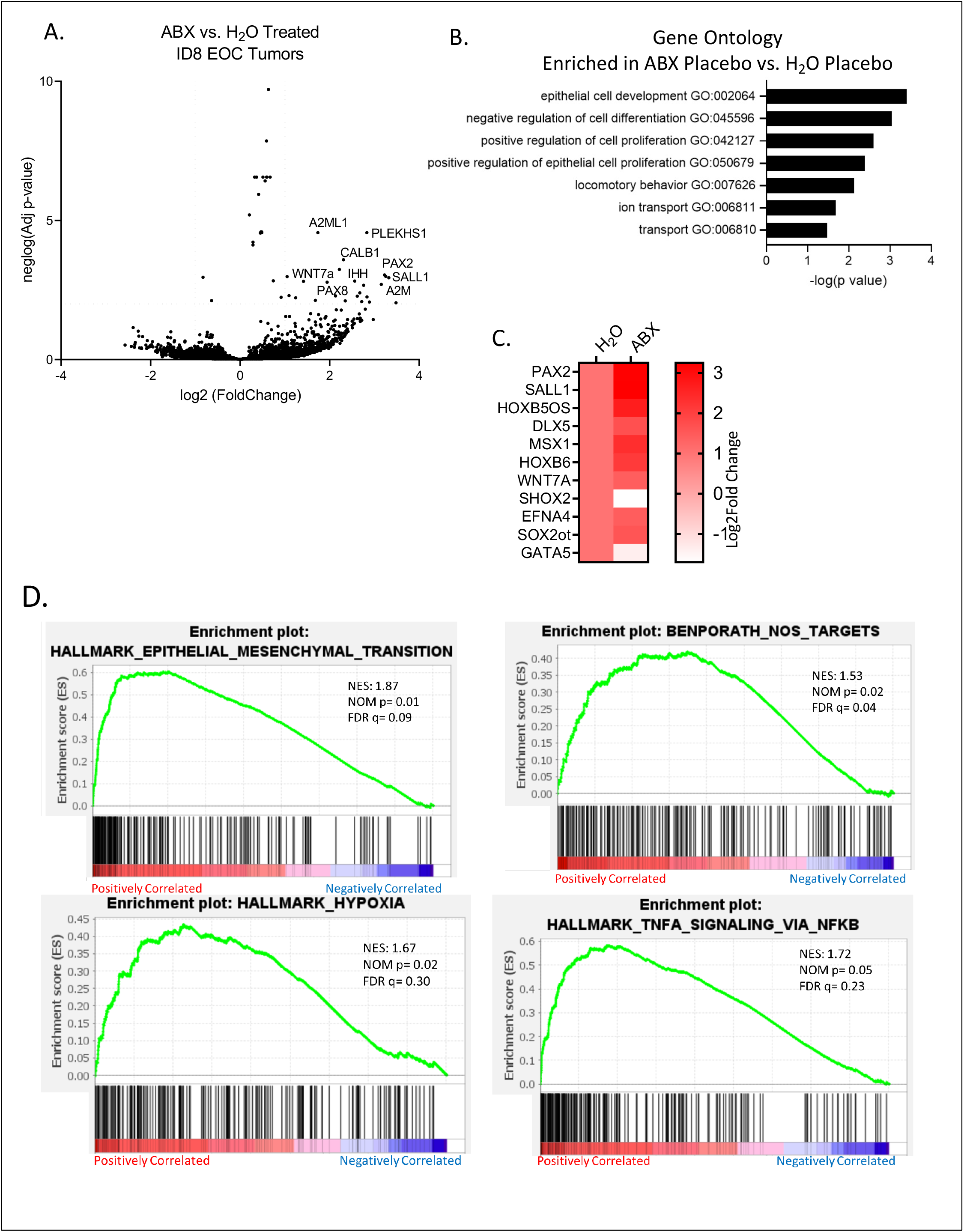
RNAseq of NSG ID8 tumors at endpoint demonstrates differentially expressed genes between treatment groups. (A) Top 100 differentially expressed genes (DEGs) ordered by p-value in the ABX Placebo group over the H_2_O Placebo group. (B) Top significant GO terms enriched in the ABX Placebo group over H_2_O Placebo group. (C) Stem cell genes enriched in the ABX Placebo group over the H_2_O Placebo group ordered by p-value. (D) Representative GSEA plots demonstrating enriched gene sets in ABX over H_2_O treated tumors.

Like the BL/6 cohort, the NSG cohort exhibited a marked relative increase in *Enterobacteriaceae* in the antibiotic treated ID8 and ID8-VEGF groups’ fecal samples, increasing more quickly in the cisplatin treated groups compared to the placebo. In addition, this cohort also saw a marked relative increase in *Paenibacillus* and *Enterococcus* (**Figure 4 A, B**).

### ABX treatment does not significantly alter ID8 or ID8-VEGF EOC tumor cell proliferation *in vitro*

To determine if tumor growth was a direct effect of ABX interaction with the EOC ID8 and ID8-VEGF cell lines, we performed tissue culture analyses. ID8 and ID8-VEGF cell lines were co-cultured with various concentrations of ABX (metronidazole, vancomycin, ampicillin and neomycin) in the same respective ratios utilized in the murine studies Following 7 days culture, there were no appreciable differences in ABX treated cell lines compared to cell lines in normal media (**Supplemental Figure 6 A, B**). Additionally, there were no changes in the determined IC50 of cisplatin on ID8 of ID8-VEGF cells following ABX treatment when compared to controls (**Supplemental Figure 6 C, D**).

**Figure 6.**
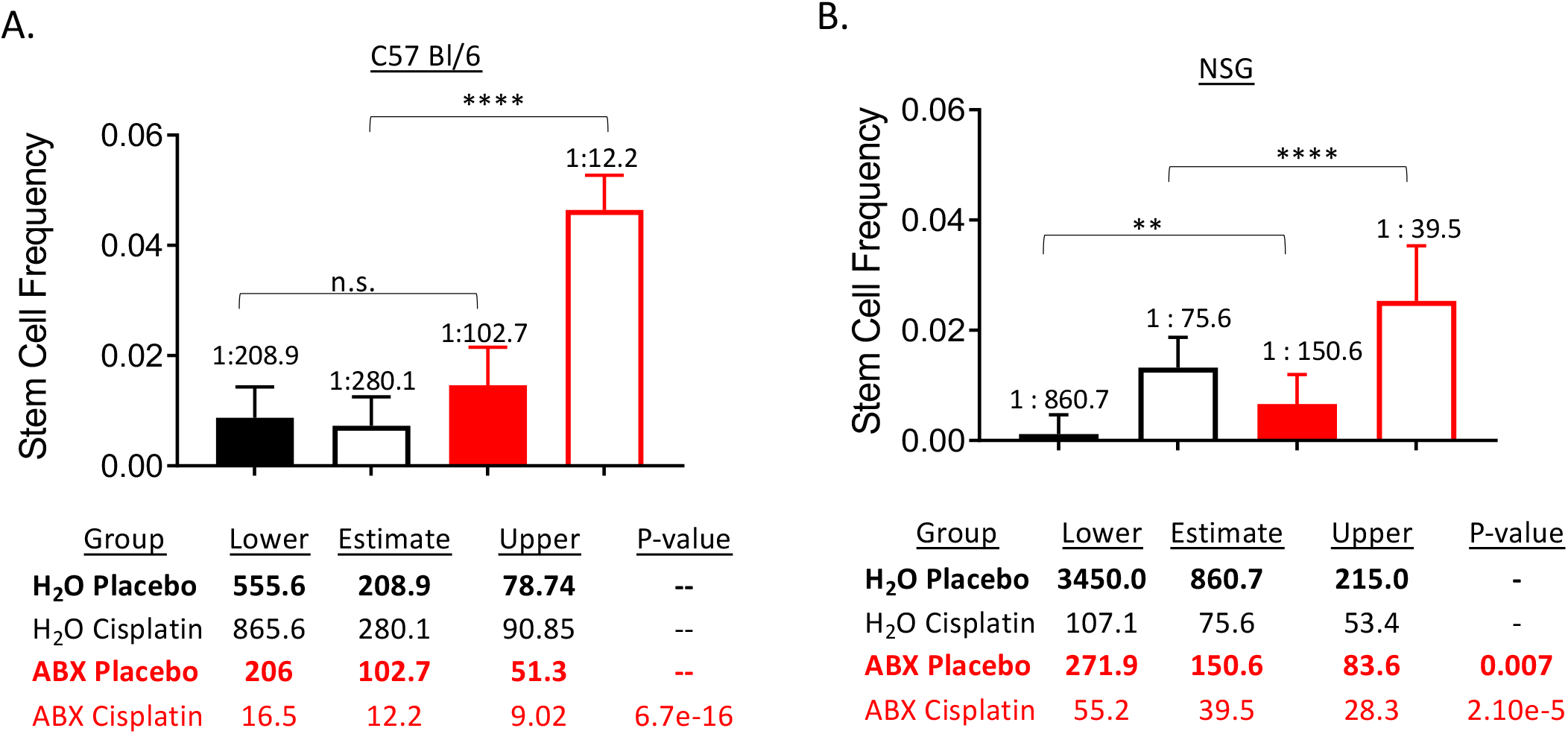
Tumor sphere initiation frequency is increased in ABX treated mice and further increased upon cisplatin therapy. ID8 tumor cells isolated from ABX treated mice had significantly increased tumor sphere initiation frequency in both BL/6 (A) and NSG (B) cohorts. **p<0.01 ***p<0.001, ***p<0.0001 n= 4 mice per group in 96 wells per condition.

### Antibiotic treatment leads to accelerated tumor growth and attenuated sensitivity to cisplatin in patient derived EOC

To ensure our observed phenotype was not unique to syngeneic EOC cell lines (ID8 and ID8-VEGF). We repeated our study paradigm in NSG mice with a human derived OV81 cell line. Following 2 weeks of ABX or control water, mice were IP injected with 5×10^6^ OV81 cells. To monitor tumor growth is this cohort of mice, cells were transduced with a luciferase reporter prior to IP injection. Following 2 weeks IP cell injection, mice were randomized into 4 groups: H_2_O Vehicle, H_2_O Cisplatin, ABX Vehicle, and ABX Cisplatin and treated as previously outlined. At endpoint, mice underwent Perkin Elmer *in Vivo* Imaging System (IVIS) imaging and total flux of photons/second was calculated to determine total tumor burden. Overall ABX therapy resulted in a significant increase in total tumor burden when normalized to initial tumor burden compared to H_2_O controls in both the vehicle and cisplatin treated groups (**Supplemental Figure 7**).

**Figure 7.**
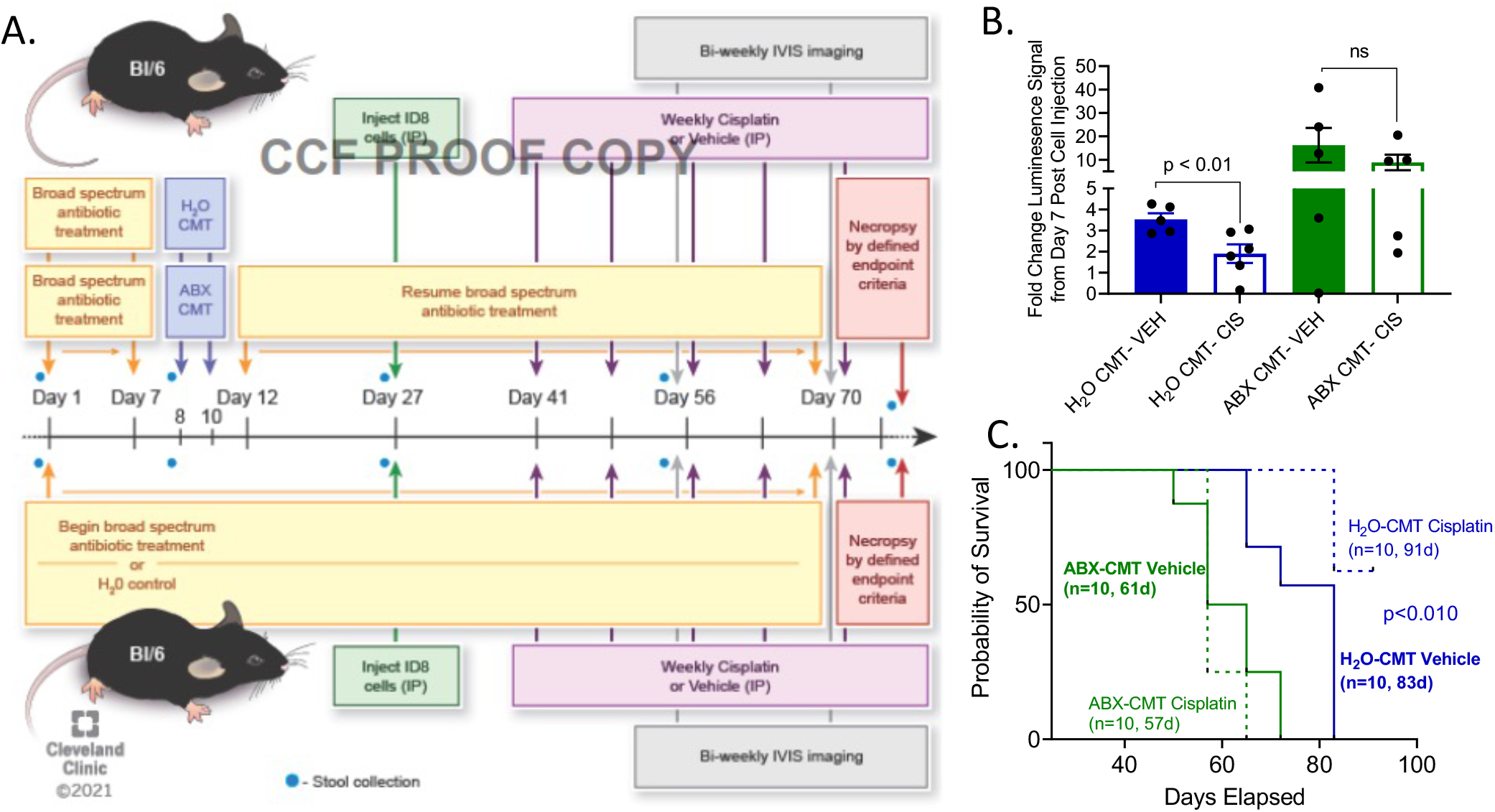
H_2_O CMT ameliorates the effects of ABX on cisplatin sensitivity and tumor progression. (A) Schematic depicting the study design for cecal microbial transplant following ABX treatment. (B) Overall fold change in tumor burden via luminescence from baseline, demonstrating an increase in cisplatin sensitivity and diminished tumor progression in H_2_O treated CMT compared to ABX treated CMT. (C) Survival analysis demonstrating diminished survival in ABX treated CMT mice regardless of cisplatin therapy compared to H_2_O treated CMT mice. n=6-8 mice per group, 2-way ANOVA.

### ABX therapy induces a stem cell phenotype in ID8 EOC tumors in NSG mice

To investigate the mechanisms of ABX disruption on EOC tumor growth and sensitivity to cisplatin, we tested for presence of bacteria in the tumors and when available in the ascites. We cultured specimen from the ABX and nonABX tumors on 10 cm bacterial plates and were not able to detect any bacterial growth. Thus, to elucidate the mechanisms underlying the impact of antibiotic therapy on tumor cells and their growth in the presence and absence of cisplatin, bulk RNAseq was performed on tumors collected from nonABX (H_2_O Placebo and H_2_O Cisplatin) and ABX (Placebo, Cisplatin) treated NSG mice at endpoint (8 weeks post ID8 cell injection). The top 100 most significantly affected genes by p-value following a pairwise comparison between the H_2_O Placebo and ABX Placebo groups (**Figure 5 A**) were analyzed utilizing DAVID software for the enrichment of GO terms. Interestingly, the most enriched GO terms included cell differentiation, proliferation, and locomotor activity in line with the increased self-renewal ability observed in the previous tumor sphere initiation assays (**Figure 5 B**).

Genes commonly associated with cancer stem cell signatures were also increased in the ABX treatment group compared to the H_2_O control group such as *SOX2, WNT7a, HOXb5, HOXb6, DLX5, MSX1, EFNA4, SALL1,* and *PAX2* genes. *SHOX2* and *GATA5* genes that promote cell differentiation were also significantly decreased (**Figure 5 C**) (63–70). Upon gene set enrichment analysis (GSEA), epithelial mesenchymal transition, NOS targets, NFkB signaling via TNFα, and hypoxia were all enriched in ABX treated tumors compared to H_2_O treated controls (**Figure 5 D**) (71–73).

As we observed increased stem cell markers based on RNAseq of tumors, we performed self-renewal assays to assess stem cell frequency in tumors from H_2_O and ABX treated mice with and without cisplatin therapy. Following endpoint necropsy, tumor tissue from each group was dissociated to single cells and plated in a limiting dilution assay. Following 14 days of incubation, sphere initiation frequency was determined indicating a significant increase in sphere initiation frequency of ID8 cells from ABX treated BL/6 mice compared to control H_2_O in both the presence and absence of cisplatin therapy (**Figure 6 A**). The observed effect on CSCs was replicated in the NSG cohorts demonstrating an apparent increase in the stem cell population following ABX therapy that is further increased in the presence of cisplatin (**Figure 6 B**).

### Cecal Microbial Transplant (CMT) from control treated mice is sufficient to inhibit tumor growth of ABX treatment on EOC tumor progression and cisplatin resistance

We tested the hypothesis that the gut microbiome of nonABX treated mice can inhibit the accelerated EOC tumor growth by performing CMT studies. Cecal content was harvested from either H_2_O or ABX treated mice following the study paradigm outlined in **Figure 7 A.** Bl/6 mice treated with ABX were orally gavaged with ABX or H_2_O cecal content, followed by IP injection with ID8 EOC cells and monitored for tumor progression with bi weekly IVIS imaging. Endpoint criteria was pre-determined as total tumor burden >150mm^3^, debilitating ascites development and/or a standard body composition score (BCS) < 2. At initial endpoint of ABX CMT treated mice, all groups were compared for total tumor progression from baseline **(Figure 7 B)**. Mice receiving H_2_O CMT had tumor burden as mice that did not receive any ABX treatment, and mice receiving ABX CMT had tumor burden similar to mice receiving ABX throughout. Survival analysis showed that ABX CMT treated mice were not responsive to cisplatin therapy with the median survival of vehicle and cisplatin treated mice at 61 and 57 days post cell injection respectively. The H_2_O CMT treated mice however, were sensitized to cisplatin therapy showing median survivals of 83 and 91 days in vehicle and cisplatin treated groups respectively **(Figure 7 C)**.

## Discussion

In recent years, there has been growing evidence demonstrating a link between the gut microbiome, carcinogenesis, and response to cancer therapy (11–18). Studies have emerged supporting, in both pre-clinical models and patient cohorts, that antibiotic therapy associated gut microbial disruption may negatively impact the efficacy of immune checkpoint inhibitors and systemic anti-cancer drugs, including platinum chemotherapy (12–17). To date, the impact of the gut microbiome upon response to chemotherapy in women with gynecologic malignancies is yet to be explored. In women diagnosed with EOC, platinum chemotherapy remains the standard treatment in primary and newly recurrent disease. Disease prognosis and treatment efficacy depends upon platinum sensitivity, with platinum resistance portending a poor prognosis with limited active treatment options. Interventions to increase platinum sensitivity and prevent development of resistance are essential to improving the care of women with EOC.

The pre-clinical studies described here demonstrate that antibiotic therapy, and concomitant changes within the gut microbiome, result in accelerated tumor growth, attenuated cisplatin sensitivity, and decreased survival. In mice who received ABX, 16S rDNA analysis demonstrated that the gut microbiome was markedly altered, compared to controls. ABX treatment led to significant reduction in diversity in the fecal microbial communities, regardless of cisplatin therapy.

We found that restoration of the gut microbiome in ABX treated mice via CMT from nonABX treated mouse cecum is sufficient to improve overall survival. This supports the conclusion that a gut derived activity/metabolite underlies the suppression of tumor growth and maintenance of chemosensitivity in EOC. We considered the alternative hypothesis that prolonged ABX treatment leads to gut microbial translocation to the peritoneum with subsequent impact on tumor growth and chemoresistance. Ascites, if available, and tumors from ABX and nonABX treated mice including cisplatin treatment conditions were cultured on BHI agar plates and incubated for 72 h under either aerobic or anaerobic conditions failed to show any bacterial growth. This analysis excludes the possibility of bacteria leaking into the peritoneum cavity due to prolonged ABX treatment and supports our conclusions of a gut derived suppressive activity in an intact microbiome.

Our findings are consistent with studies investigating the impact of the gut microbiome upon response to platinum chemotherapy and immune checkpoint inhibitor treatment for non-gynecologic cancers. Indeed, Routy et al, showed resistance to immune checkpoint inhibitors was linked to abnormal gut microbiome composition (18). Following disruption of the gut microbiome through ABX treatment, response to CpG-oligonucleotide immunotherapy and cyclophosphamide was impaired secondary to reduced cytokine production, lower production of reactive oxygen species and diminished cytotoxicity. In a separate study, Iida et al., demonstrated that disruption of the gut microbiome through ABX, impaired platinum response, with decreased tumor regression and survival, in animal models of colon cancer and lymphoma (14). Notably, the studies of Cho and colleagues (74) indicate that disruption of the gut microbiome with ABX is sufficient to delay induced spontaneous ovarian cancer development. In their studies, Cheng, et al. used a genetically engineered mouse model to induce ovarian cancer in presence and absence of ABX treatment. While the data suggest a disruption of the microbiome leads to reduced tumors in contrast to our findings, the experimental paradigm underlying tumor development and progression are very different. Indeed, the conclusions of Cheng et al. indicate changes in the microbiome composition can affect ovarian cancer growth and complement our findings.

Similar findings to ours, have been documented in cohorts of patients undergoing platinum chemotherapy for non-gynecologic cancers. Among 800 patients enrolled on clinical trials receiving cyclophosphamide or cisplatin for chronic lymphocytic leukemia or lymphoma, receipt of ABX targeting gram positive species during chemotherapy was associated with significantly decreased treatment response, time to recurrence and survival (17). These findings suggest that presence of specific bacterial populations may be essential for treatment response. In our studies, we identified significantly higher relative abundance of potentially pathogenic Gram-negative *Enterobacteriaceae* and *Parasutterella* species. Notably, among animals treated with cisplatin, these bacteria were further increased, in the absence of facultative Gram-positive bacteria, to those who received placebo.

Although not yet studied in relation to cancer, an increase in the abundance of *Parasutterella* has been associated with dysbiosis of the gut microbiome and alterations in liver metabolism (61, 62, 75).

In NSG mice, we determined that disruption of the immune system does not directly contribute to increased tumor growth or reduced platinum sensitivity in the setting of microbial disruption. In addition, no differences in myeloid and lymphoid populations were observed in the ascetic fluid of mice following antibiotic treatment, compared to controls. Our results identify that in the setting of an altered microbiome, response to platinum chemotherapy is reduced and populations of chemo-resistant cancer stem cells are increased based on tumorsphere initiation frequency and transcriptional reprogramming to an increased cancer stem cell-like state.

The relationship between enhanced populations of cancer stem cells and platinum resistance in EOC is well-documented in the literature (54, 69, 72, 76-79). Specifically, we identified *SOX2, WNT7a, HOXb5, HOXb6, DLX5, MSX1, EFNA4, SALL1,* and *PAX2* as being significantly upregulated in ID8 tumors of the ABX group over the control H_2_O group. Many of these genes are markers of pluripotency and regulation as well as markers of long-term stemness (63, 64, 66-69). SALL1 interacts with *NANOG* a well-established pluripotency transcription factor, to suppress differentiation (68). Conversely, *SHOX2* and *GATA5* genes that favor differentiation, were significantly decreased providing further evidence that ID8 tumor cells from ABX mice are more stem like and undifferentiated than ID8 tumor cells from H_2_O treated mice (65, 70, 80). Upon KEGG pathway analysis of enriched genes in the ABX group, multiple signaling pathways including P13K/AKT/mTOR and WNT, were identified and currently under investigation as targets for ovarian cancer therapies (73, 81). Further investigation is needed to understand the mechanism through which the gut microbiome interacts with cancer stem cells and how this specifically drives platinum resistance in EOC.

The clinical implications of the gut microbiome impacting platinum response in EOC are significant. Primarily, antibiotic therapy is often unavoidable in the care of patients with EOC following cyto-reductive surgery or during chemotherapy. However, here the evidence supports the need for judicious selection of antibiotics and duration of dosing as changes to the gut microbiome may negatively impact oncologic outcomes. It also supports the concept that measures to prevent infection in women with EOC, and therefore avoid antibiotic treatments, should be prioritized. Most importantly, understanding that disruption of gut microbiota impacts EOC growth and platinum chemotherapy introduces the potential for development of targeted therapeutics, designed to restore an intact gut microbiome, which may represent promising strategies to treat EOC and combat platinum resistance in the future.

## Conclusions

We identify a role for the disruption of gut microbiota in enhanced tumor growth and reduced sensitivity to platinum chemotherapy in pre-clinical models of EOC. Further investigation is critically needed to understand how the tumor microenvironment, and cancer stem cells specifically, communicate with the gut microbiome to drive platinum resistance in EOC. Answers to these questions will provide important insights as to whether microbe directed interventions to the gut microbiome can be used to impact the response to platinum chemotherapy.

## Supporting information

Chambers Supp Data

## Abbreviations

EOC: epithelial ovarian cancer
ABX: antibiotic
CMT: cecal microbiome transplant
C57BL/6J: BL/6
TAUS: transabdominal ultrasound
NSG: NOD.Cg-Prkdc<scid>Il2rg<tm1Wjl>SzJ
AVS: amplified sequence variants
bps: base pairs
GSEA: gene set enrichment analysis
CSC: cancer stem cell
BCS: body composition score
IVIS: in vivo imaging system
DADA: divisive amplicon denoising algorithm
PCoA: principal coordinate analysis
MDS: multidimensional scaling
PERMANOVA: permutational multivariate analysis of variance
FDR: false discovery rate

## Declarations

### Ethics approval and consent to participate

Not Applicable

### Consent for publication

All authors have consented to publication.

### Availability of data and material

All tumor RNAseq and 16S rRNA sequencing data is publicly available upon acceptance of publication.

### Authors’ contributions

OR, LC, RV, EE, MD, PA, JC, JL, and JMB conceived and designed the studies. OR, CM, RV, PGR provided administrative support and funding. LC, EE, LT, CB, DB, AMH, and ZA performed the mouse studies including necropsy, tissue harvest, in vitro studies, and assembly of data. LC, EE, OR, CM, PA, JC, DB, and AJP performed the data analysis and interpretation. NS, PB, and EE performed the 16S rDNA and RNAseq analysis. EE, LC, OR, CM, NS, PB, AMH, JL, and JMB participated in manuscript writing. All authors read, edited, and approved the final approval of the manuscript.

## Acknowledgements

The authors would like to acknowledge members of the Reizes laboratory for their collaborative effort and insight toward the completion of the studies within this manuscript, as well as the LRI Core Facilities that played a role in data collection, analysis, or interpretation of the findings presented within the manuscript including Microbiome Analytics and Composition Core Facility, Image Core, Histology Core, and Microbial Culture and Engineering Core. DB received support from NIH F32 CA243314. Dr. Reizes is the Laura J. Fogarty Endowed Chair in Uterine Cancer Research. Research in the Reizes laboratory is funded by VeloSano Bike to Cure, Center of Research Excellence in Gynecologic Cancer, and the Department of Defense.

## Supplemental Material

**Supplemental Figure 1. Representative multi –parameter gating strategy for myeloid and lymphoid cell populations**.

**Supplemental Figure 2. Myeloid and lymphoid immune cell population analysis from ascites of Bl/6 and NSG ID8 VEGF tumor bearing mice at endpoint.** Following ABX therapy, BL/6 ID8 VEGF ascites did not exhibit significant alterations in myeloid (A) or lymphoid populations (B) by flow cytometry analysis. NSG ID8 VEGF ascites did not exhibit significant alterations in myeloid (C) populations. Values are represented as percent (%) among CD45+ staining. BL/6 n=4 mice per group, NSG n=5 mice per group

**Supplemental Figure 3.** Myeloid and lymphoid immune cell population analysis from Bl/6 ID8 tumor-bearing splenocytes and bone marrow. Following ABX therapy, BL/6 ID8 splenocytes (A and B) or bone marrow (C and D) did not exhibit significant alterations in myeloid or lymphoid cell populations by flow cytometry analysis. Values are represented as percent (%) among CD45+ staining. n=3 mice per group, error bars represent SEM.

**Supplemental Figure 4. BL/6 16S Alpha and Beta Diversity.** A. Alpha diversity of Bl/6 ID8 stool 16S over time. B. Alpha diversity of Bl/6 ID8-VEGF stool 16S over time. C. Beta diversity of Bl/6 ID8 and ID8-VEGF over time. n= 8 mice per group, PERMANOVA

**Supplemental Figure 5. NSG 16S Alpha and Beta Diversity.** A. Alpha diversity of NSG ID8 stool 16S over time. B. Alpha diversity of NSG ID8-VEGF stool 16S over time. C. Beta diversity of NSG ID8 and ID8-VEGF over time. n= 8 mice per group, PERMANOVA

**Supplemental Figure 6.** ABX treatment does not significantly alter tumor cell proliferation or cisplatin sensitivity *in vitro.* Following 4 days co-culture of ID8 (A) and ID8 VEGF (B) with ABX at vary concentrations, no significant change in proliferation was observed compared to cells cultured with media alone. The IC50 of Cisplatin was not significantly altered when co cultured with ABX compared to control media in ID8 (C) or ID8 VEGF (D) cells in vitro. n= 3 replicates per data point, representative of 3 biological replicates.

**Supplemental Figure 7. ABX treatment decreases sensitivity of OV81 cells to cisplatin therapy, resulting in increased tumor growth.** Mice treated with antibiotics exhibited increased overall OV81 tumor growth in both the presence and absence of cisplatin therapy compared to the H_2_O treated control. ***p<0.001, student’s t test.

**Supplemental Figure 8. Survival was enhanced with cisplatin treatment in H_2_O and H_2_O-CMT but not in ABX and ABX-CMT treated groups.** A. Survival plots of H_2_O treated groups demonstrating the similarity in survival between H_2_O CMT (blue) and H_2_O (black) vehicle (solid lines) and H_2_O CMT (blue) and H_2_O (black) cisplatin (dashed lines) groups. B. Survival plots of ABX treated groups demonstrating no significant advantage of cisplatin therapy in ABX-CMT (green) or ABX (red) groups, and a significant decrease in survival in ABX-CMT (green) compared to ABX (red) cisplatin treated groups (dashed lines).

**Supplemental Table 1. Myeloid and lymphoid antibody lists utilized for ascites, splenocyte and bone marrow analysis.**

