## Supplementary material for "Disruption of the gut microbiota attenuates epithelial ovarian cancer sensitivity to cisplatin therapy": Chambers Supp Data

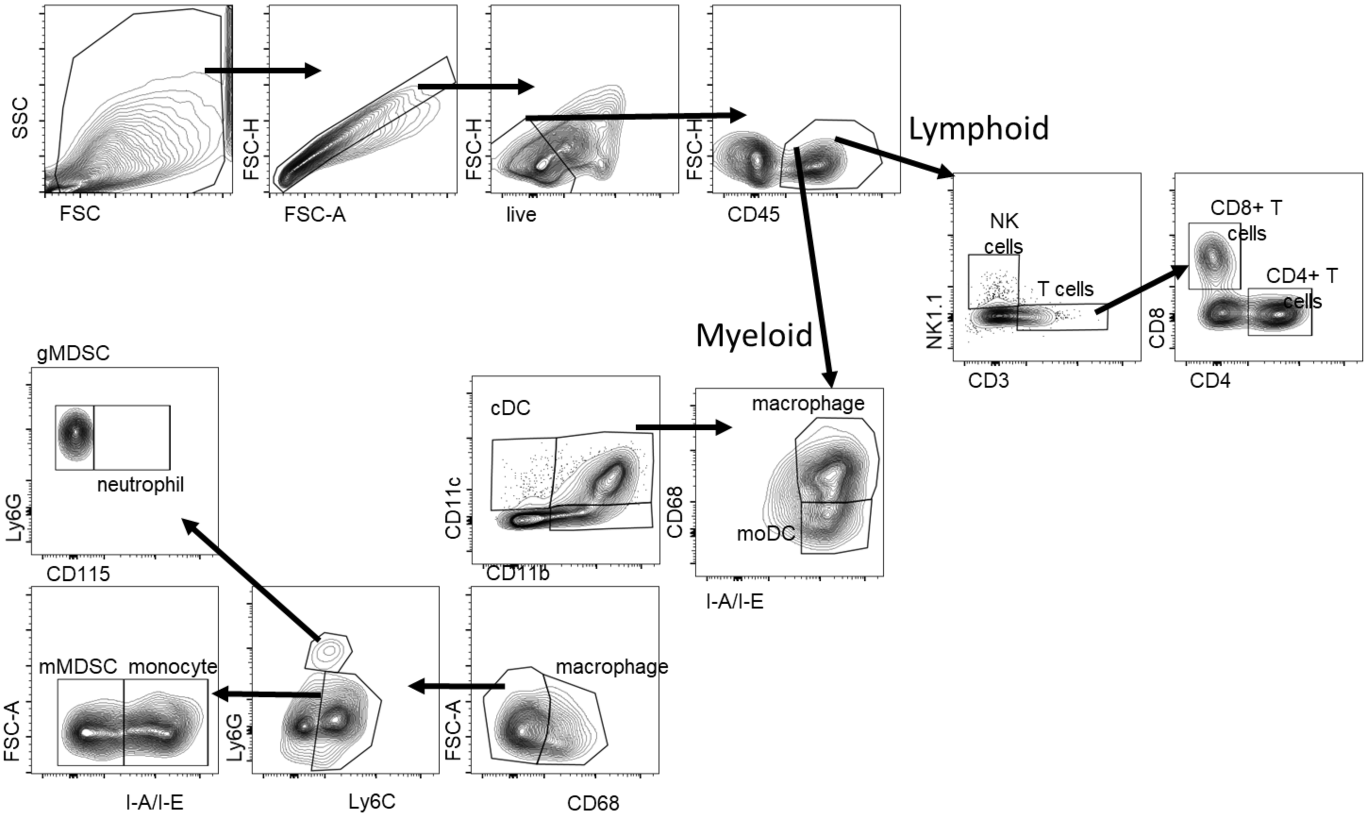


**Supplemental Figure 1. Representative multi –parameter gating strategy for myeloid and lymphoid cell populations.**

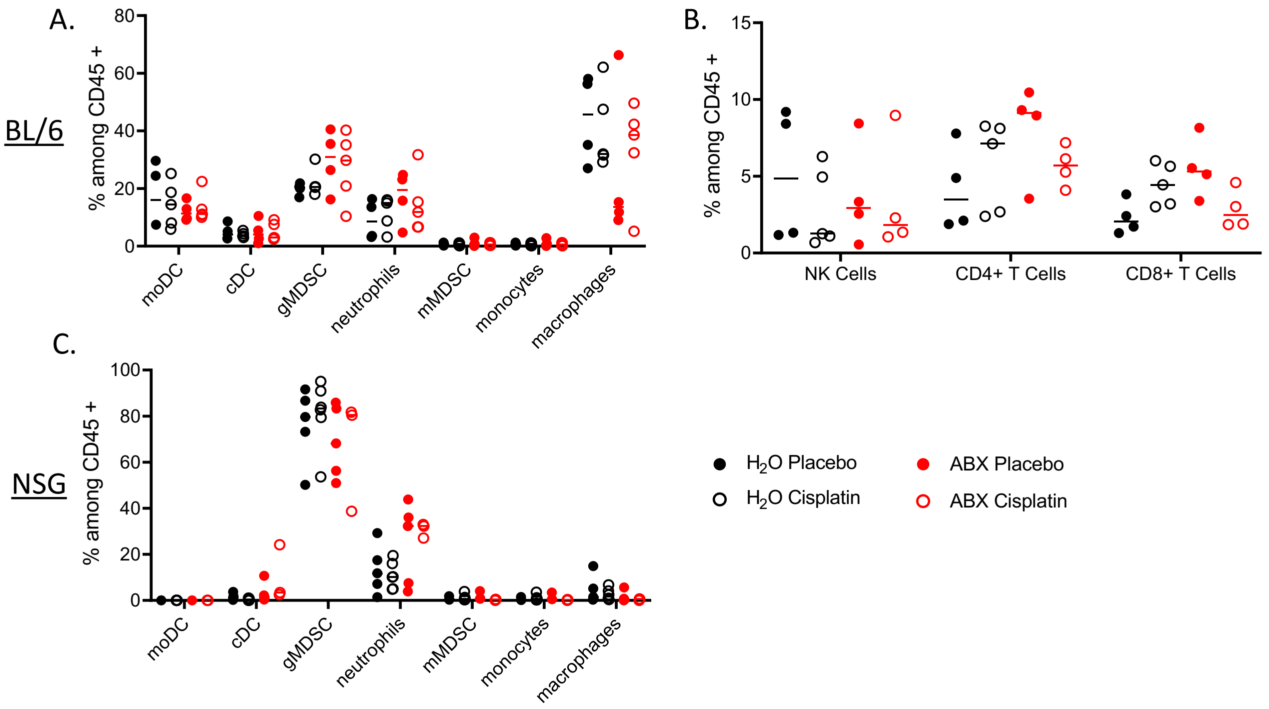


**Supplemental Figure 2. Myeloid and lymphoid immune cell population analysis from ascites of Bl/6 and NSG ID8 VEGF tumor bearing mice at endpoint.** Following ABX therapy, BL/6 ID8 VEGF ascites did not exhibit significant alterations in myeloid (A) or lymphoid populations (B) by flow cytometry analysis. NSG ID8 VEGF ascites did not exhibit significant alterations in myeloid (C) populations. Values are represented as percent (%) among CD45+ staining. BL/6 n=4 mice per group, NSG n=5 mice per group

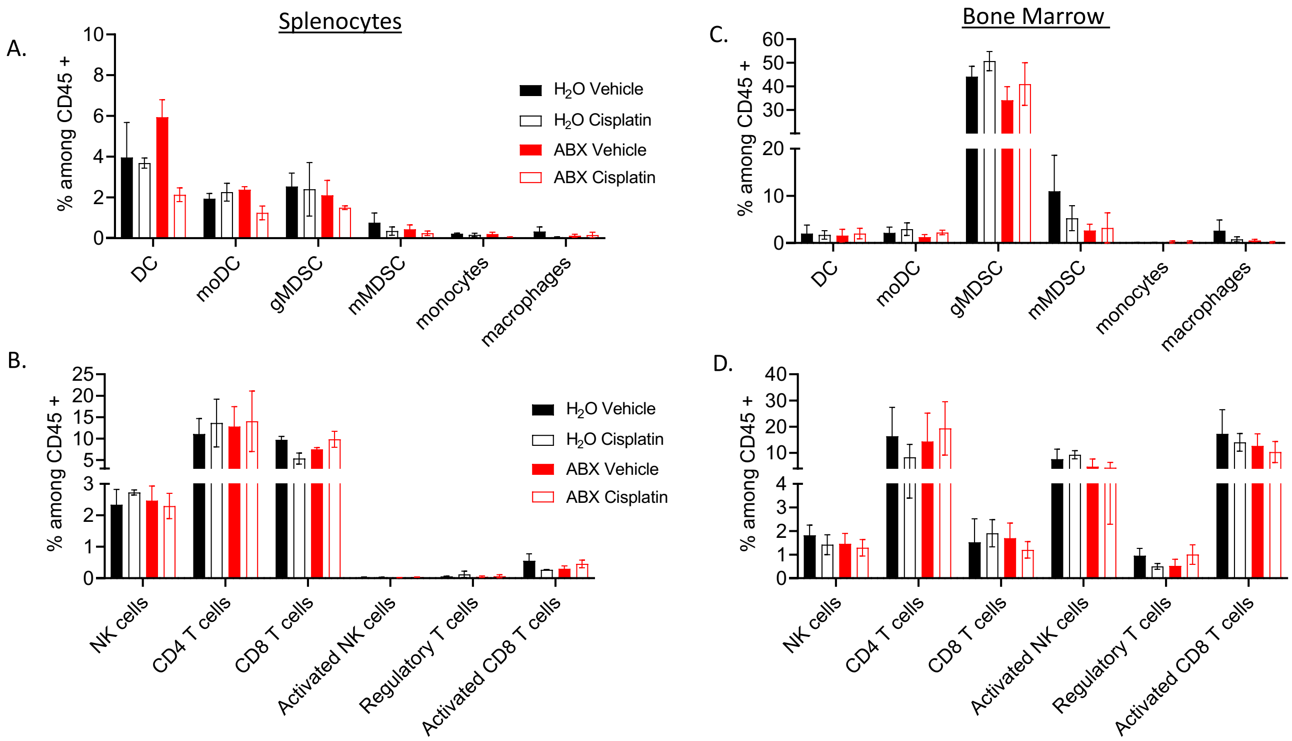


**Supplemental Figure 3.** Myeloid and lymphoid immune cell population analysis from Bl/6 ID8 tumor-bearing splenocytes and bone marrow. Following ABX therapy, BL/6 ID8 splenocytes (A and B) or bone marrow (C and D) did not exhibit significant alterations in myeloid or lymphoid cell populations by flow cytometry analysis. Values are represented as percent (%) among CD45+ staining. n=3 mice per group, error bars represent SEM.

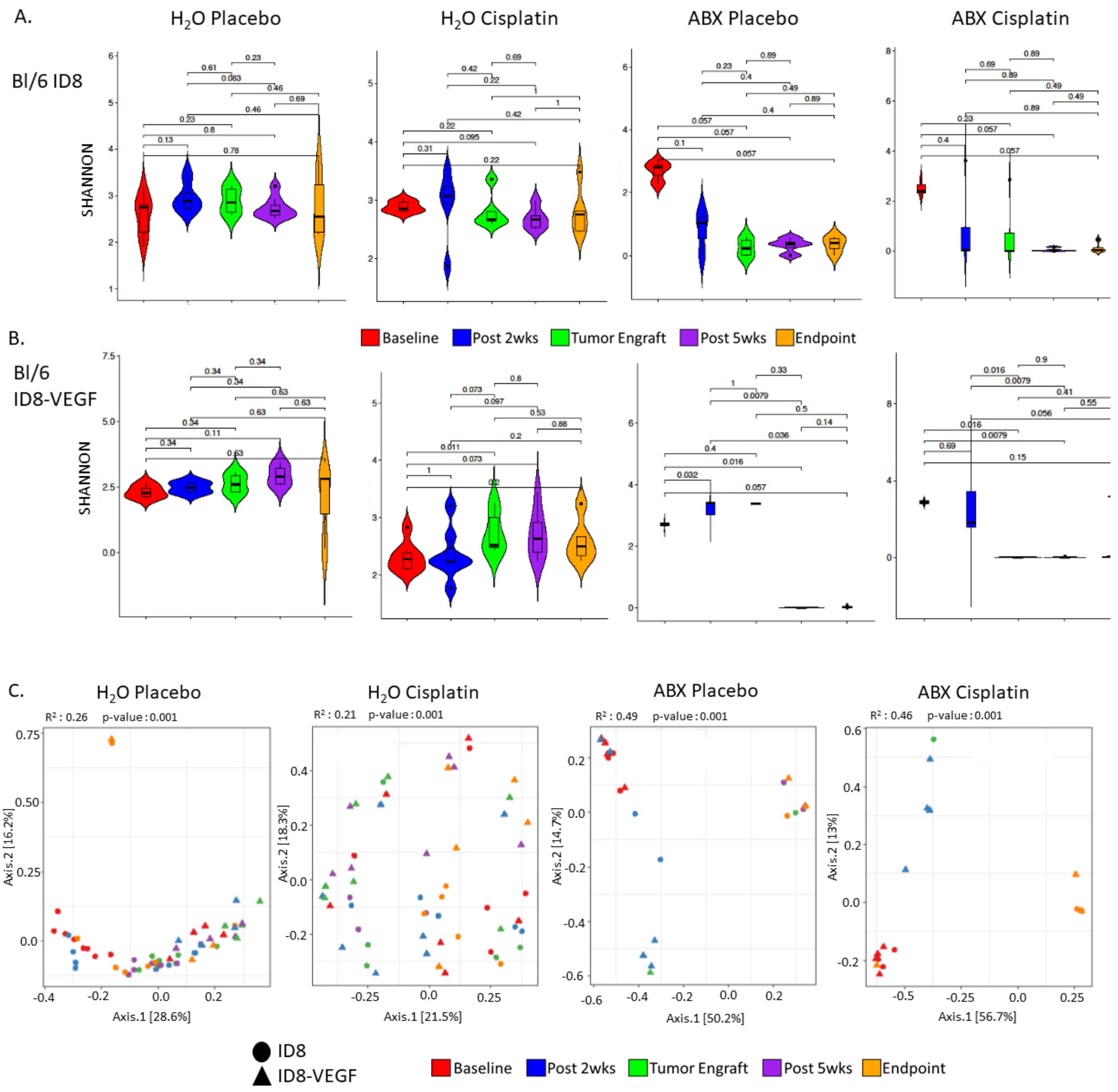


**Supplemental Figure 4. BL/6 16S Alpha and Beta Diversity.** A. Alpha diversity of Bl/6 ID8 stool 16S over time. B. Alpha diversity of Bl/6 ID8-VEGF stool 16S over time. C. Beta diversity of Bl/6 ID8 and ID8-VEGF over time. n= 8 mice per group, PERMANOVA

Supplemental Figure 5

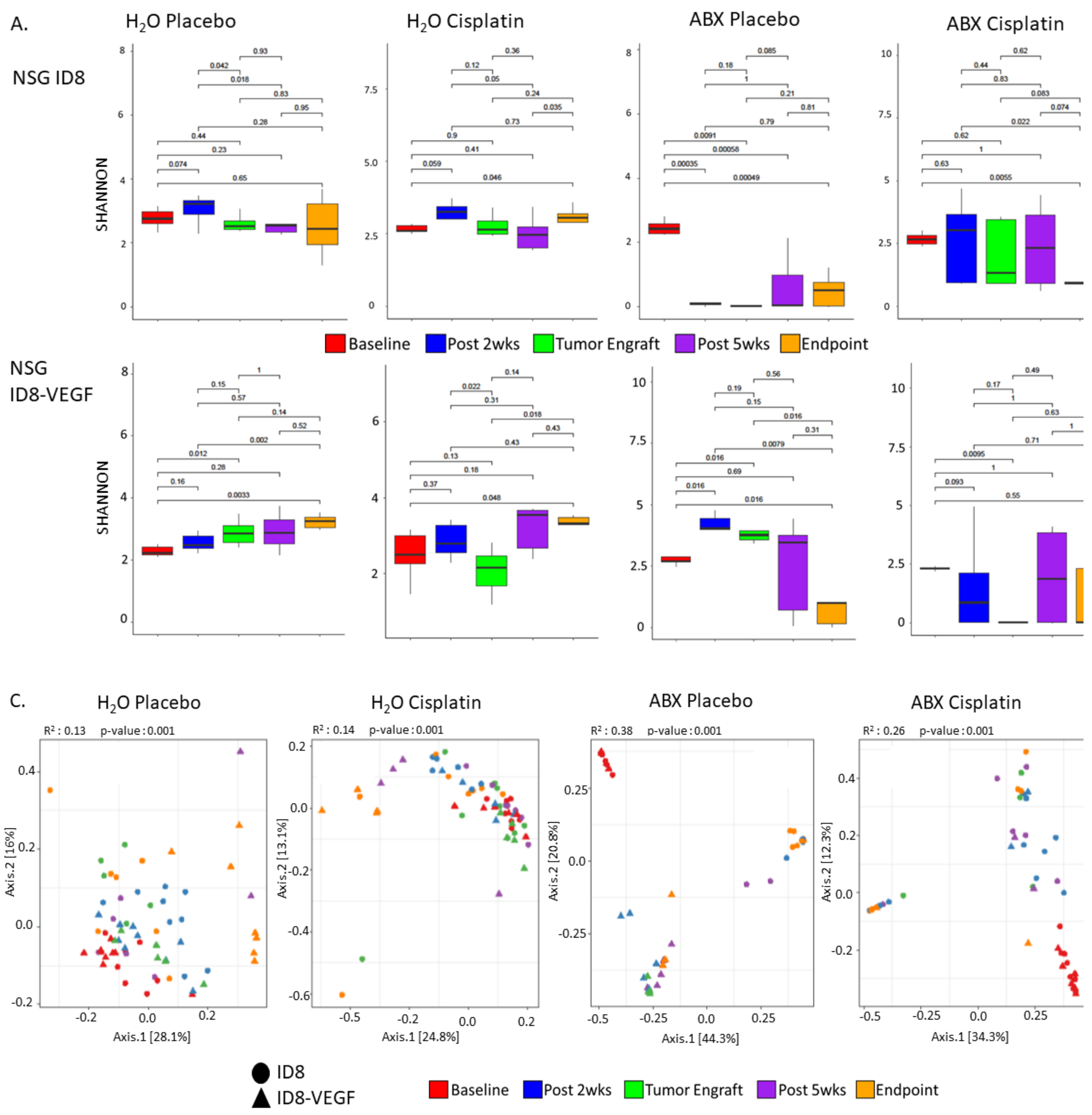


**Supplemental Figure 5. NSG 16S Alpha and Beta Diversity.** A. Alpha diversity of NSG ID8 stool 16S over time. B. Alpha diversity of NSG ID8-VEGF stool 16S over time. C. Beta diversity of NSG ID8 and ID8-VEGF over time. n= 8 mice per group, PERMANOVA

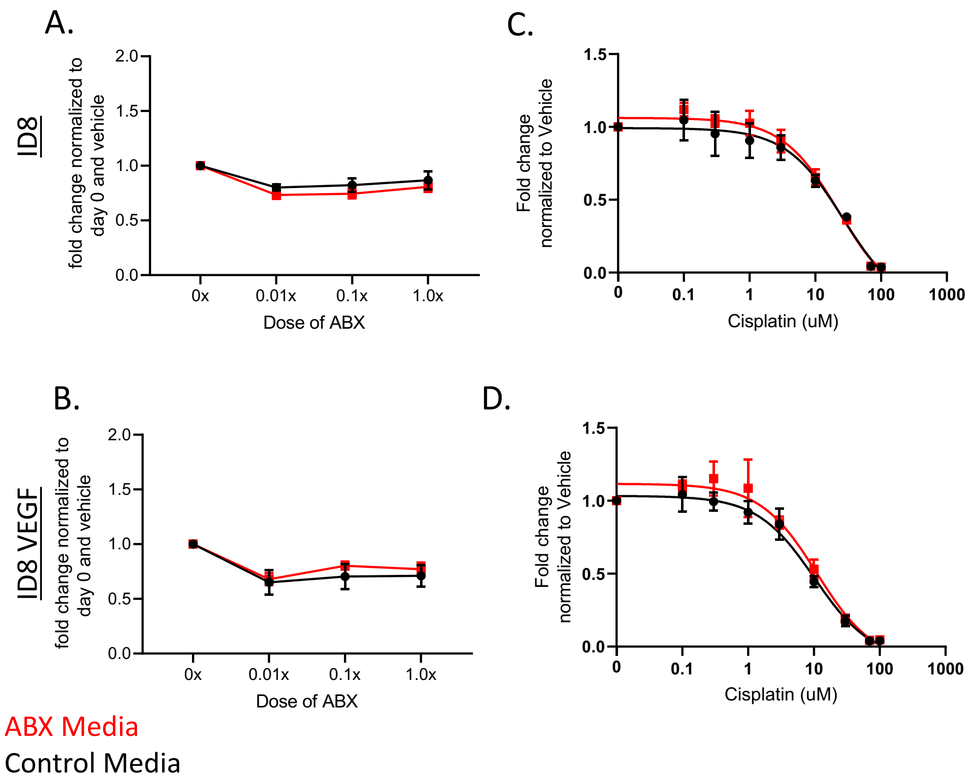


**Supplemental Figure 6.** ABX treatment does not significantly alter tumor cell proliferation or cisplatin sensitivity *in vitro.* Following 4 days co-culture of ID8 (A) and ID8 VEGF (B) with ABX at vary concentrations, no significant change in proliferation was observed compared to cells cultured with media alone. The IC50 of Cisplatin was not significantly altered when co cultured with ABX compared to control media in ID8 (C) or ID8 VEGF (D) cells in vitro. n= 3 replicates per data point, representative of 3 biological replicates.

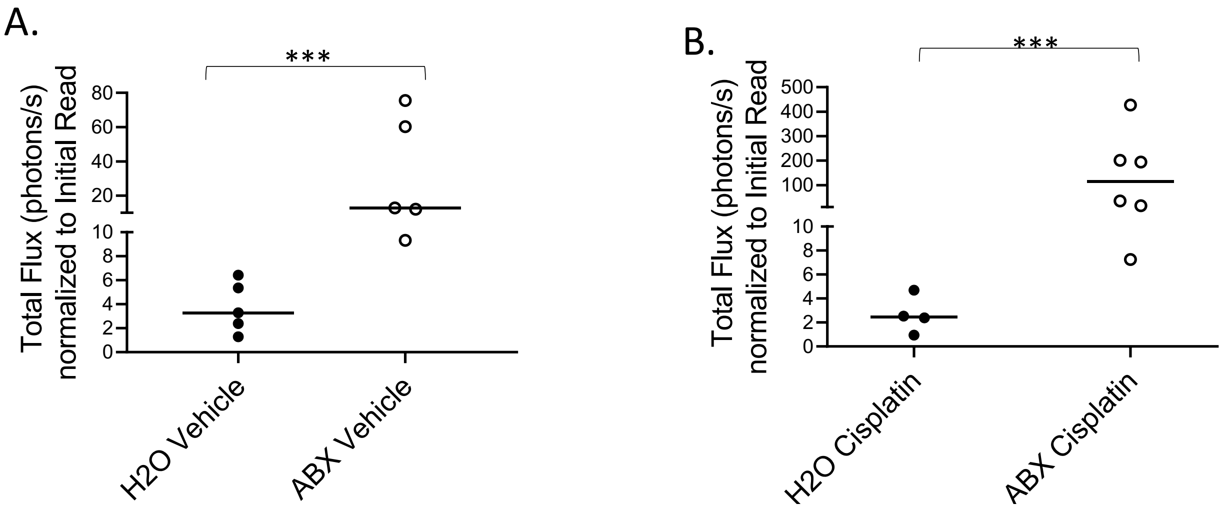


**Supplemental Figure 7. ABX treatment decreases sensitivity of OV81 cells to cisplatin therapy, resulting in increased tumor growth.** Mice treated with antibiotics exhibited increased overall OV81 tumor growth in both the presence and absence of cisplatin therapy compared to the H2O treated control. ***p<0.001, student's t test.

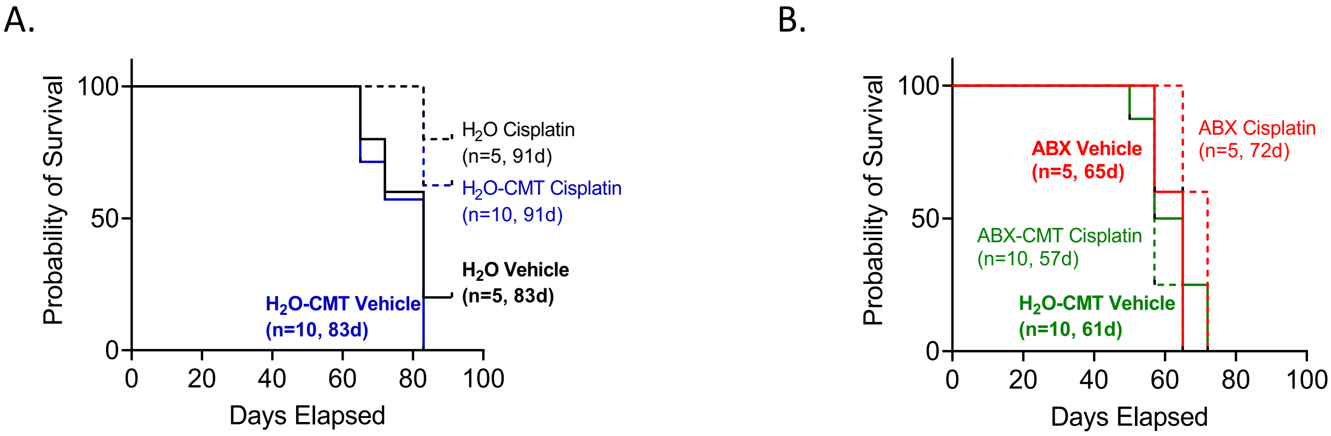


**Supplemental Figure 8. Survival was enhanced with cisplatin treatment in $H_2O$ and $H_2O$-CMT but not in ABX and ABX-CMT treated groups.** A. Survival plots of $H_2O$ treated groups demonstrating the similarity in survival between $H_2O$ CMT (blue) and $H_2O$ (black) vehicle (solid lines) and $H_2O$ CMT (blue) and $H_2O$ (black) cisplatin (dashed lines) groups. B. Survival plots of ABX treated groups demonstrating no significant advantage of cisplatin therapy in ABX-CMT (green) or ABX (red) groups, and a significant decrease in survival in ABX-CMT (green) compared to ABX (red) cisplatin treated groups (dashed lines).

| Myeloid Panel | | |
|---|---|---|
| **Target** | **Fluorophore** | **Catalog Number** |
| CD45 | PerCP/Cy5.5 | 103132 |
| CD11b | APC | 101211 |
| Ly6-C | AF700 | 128024 |
| Ly6-G | PE Cy7 | 127618 |
| CD11c | BV421 | 117330 |
| CD68 | APC Cy7 | 137024 |
| CD206 | PE | 141706 |
| CD115 | BV605 | 121433 |
| IA/IE | FITC | 107606 |
| **Lymphoid Panel** | | |
| **Target** | **Fluorophore** | **Catalog Number** |
| CD45 | PerCP/Cy5.5 | 103132 |
| CD69 | AF700 | 104539 |
| NK1.1 | BV421 | 108741 |
| CD25 | BV605 | 102035 |
| CD279 | FITC | 135214 |
| CD4 | PE Cy7 | 25-0041-82 eBioscience |
| CD8 | APC | 100712 |
| CD3 | APC Cy7 | 100330 |

**Supplemental Table 1. Myeloid and lymphoid antibody lists utilized for ascites, splenocyte and bone marrow analysis.**
